## Supplemental Information for "Protein structural features predict responsiveness to pharmacological chaperone treatment for three lysosomal storage disorders"

Supplemental Figures

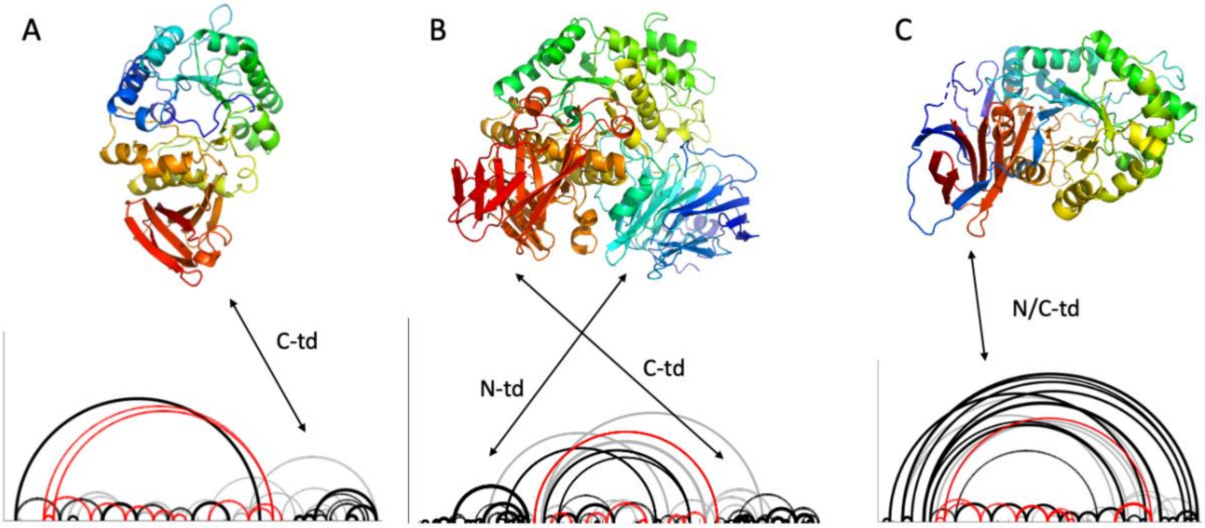


**Fig S1. Diagrams of proteins with mutants associated with lysosomal storage disorders.** (A) alpha-galactosidase A (Fabry disease), (B) acid alpha-glucosidase (Pompe disease) (C) glucocerebrosidase (Gaucher disease). Topology diagrams are shown below the protein representations, where curves connect contacting secondary structural elements along the chain, with contacts between beta strands in black, contacts between alpha helices in. red, and other contacts in gray.

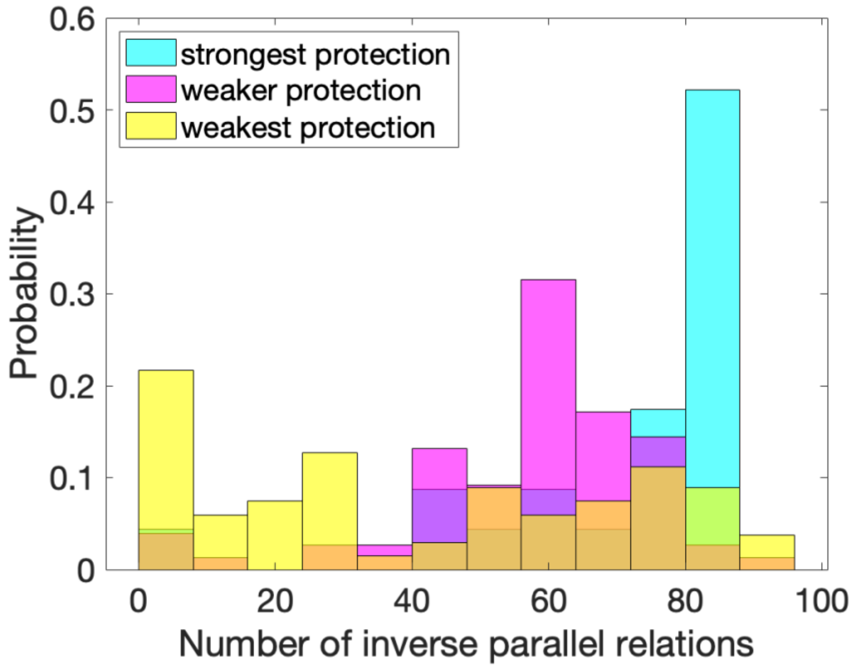


**Fig S2. Number of inverse parallel relations, by protection from exchange in mass spectrometry experiments.** Crystal structure 1J5T is referenced. Strongest protection refers to residues in β4α4. Weaker protection is conferred in α1β2 and β5α5β6α6β7. Weakest protection refers to all other residues. T-test p values are: 0.005 for strongest and weaker protection, $6x10^{-5}$ for strongest and weakest protection, and $2x10^{-4}$ for weaker and weakest protection.

Supplemental Tables

**Table S1. Optimization of Fabry decision tree complexity**

| complexity parameter | 0.01 | 0.012 | 0.014 | 0.016 | **0.018** | 0.02 | 0.022 | 0.024 | 0.026 |
|---|---|---|---|---|---|---|---|---|---|
| MCC Pompe | 0.412 | 0.443 | 0.443 | 0.443 | **0.492** | 0.492 | 0.492 | 0.492 | 0.36 |
| MCC Fabry | 0.513 | 0.449 | 0.449 | 0.449 | **0.393** | 0.376 | 0.376 | 0.376 | 0.323 |
| Nodes | 14 | 8 | 8 | 8 | **5** | 4 | 4 | 4 | 2 |

**Table S2. Leading auto-ML models**

| Dataset | Best performing model |
|---|---|
| Fabry | StackedEnsemble_BestOfFamily_AutoML_20210422_151045 |
| Pompe | XGBoost_grid__1_AutoML_20210422_152715_model_28 |
